# Plant hydraulics coordinated with photosynthetic traits and climate

**DOI:** 10.1101/2021.03.02.433324

**Authors:** Huiying Xu, Han Wang, I. Colin Prentice, Sandy P. Harrison, Ian J. Wright

## Abstract

The coupling between water loss and carbon dioxide uptake drives the coordination of plant hydraulic and photosynthetic traits. Analysing multi-species measurements on a 3000 m elevation gradient, we found that hydraulic and leaf-economic traits were less plastic, and more closely associated with phylogeny, than photosynthetic traits. The two trait sets are linked by the sapwood-to-leaf area ratio (Huber value, *v*_H_), shown here to be codetermined by sapwood hydraulic conductance (*K*_S_), leaf mass-per-area (LMA) and photosynthetic capacity (*V*_cmax_). Substantial hydraulic diversity was related to the trade-off between *K*_S_ and *v*_H_. Leaf drought tolerance (inferred from turgor loss point, –π_tlp_) increased with wood density, but the trade-off between hydraulic efficiency (*K*_S_) and –π_tlp_ was weak. The least-cost optimality framework was extended to predict trait (*K*_S_-dominated) and environmental (temperature-dominated) effects on *v*_H_. These results suggest an approach to include photosynthetic-hydraulic coordination in land-surface models; however, prediction of non-plastic trait distributions remains a challenge.

---

Water transport is essential for plant survival and growth. Hydraulic failure triggers death under severe drought ^1^, and differences in hydraulic traits can be used to predict drought-induced tree mortality ^2^. Photosynthesis is constrained by hydraulics because water transported through the xylem must replenish water lost through stomata during CO_2_ uptake ^3^. Empirical studies ^4–6^ and optimality arguments ^7^ support a tight coordination between hydraulic and photosynthetic traits. Nonetheless, quantitative understanding of their relationships remains incomplete ^8^. Embedding plant hydraulics in vegetation and land-surface models is desirable ^9,10^, not least because an improved understanding of drought effects on photosynthesis and transpiration could remove a leading source of uncertainty in global models ^11^. This situation provides strong motivation for theoretical and empirical research on how whole-plant hydraulic traits are related to (better-studied) leaf photosynthetic traits.

The ratio of sapwood area to subtended leaf area (the Huber value, *v*_H_) links whole-plant to leaf processes ^8,12^. All else equal, sapwood area determines the maximum water supply rate, while *v*_H_ in turn depends on carbon allocation to stems versus leaves. Plants with low *v*_H_ tend to have low leaf mass-per-area (LMA) and low leaf stable carbon isotope ratios (δ^13^C), implying a high ratio of leaf-internal to ambient CO_2_ (χ); high maximum CO_2_ assimilation rate (*A*_sat_); high leaf water potential at the turgor loss point (π_tlp_, a negative quantity); and high sapwood-specific hydraulic conductance (*K*_S_) _6,8,12_. Previous studies have also shown that hydraulic traits are influenced by environmental variables, particularly aridity ^13–16^, in a coordinated way. Drought-adapted plants are characterized by reduced water supply through stems (low hydraulic efficiency, *K*_S_; e.g. associated with narrow conduits) and/or reduced demand (high *v*_H_), and increased leaf hydraulic safety (low π_tlp_). Photosynthetic and leaf-economic traits are also influenced by climate. χ increases with growth temperature, and decreases with vapour pressure deficit (*D*) and elevation ^17,18^. Photosynthetic capacity (maximum Rubisco carboxylation rate, *V*_cmax_) increases with light, and weakly with temperature and VPD ^19^. LMA increases with light and aridity, and decreases with temperature ^20,21^.

Optimality theory allows testable predictions about trait-trait coordination and can also provide strong explanations for observed responses of traits to environment^22^. Among photosynthetic traits, analyses of δ^13^C data have shown quantitative agreement between observed and theoretically predicted environmental responses of χ ^17,18,23^. Smith, et al. ^19^, similarly, used optimality theory to predict the observed environmental responses of *V*_cmax_ in a global data set. Sperry, et al. ^24^ integrated hydraulic traits with a photosynthesis model to predict stomatal conductance using optimality theory. Less attention has been paid to applying optimality theory to predict leaf-economic or hydraulic traits. Here we investigate the relationships among photosynthetic, leaf-economic and hydraulic traits, and between these traits and climate, using field data collected from 11 sites in the Gongga Mountain region of western China. We extend the optimality framework of Prentice, et al. ^17^ and Wang, et al. ^18^, which hypothesizes that plants minimize the total cost of maintaining the capacities for photosynthesis and water transport, in order to make explicit quantitative predictions of these relationships. *K*_S_ and *V*_cmax_ are two key traits related to water transport and photosynthesis/water demand, respectively. Based on the requirement that water transport through xylem must equal water loss via stomata, our analysis indicates a key role for *v*_H_ in achieving this requirement ^8^ and a positive relationship between *v*_H_ and *V*_cmax_ but a negative one with *K*_S_. Our model provides a new theoretical basis to understand the variations of *v*_H_ along environmental gradients.

## Results

The Gongga Mountain region of Sichuan Province, China (SI Fig. 1) extends from 29° 22’ to 29° 55’ N and 101° 1’ to 102° 9’ E and spans an elevation range from near sea level to 8000 m. This elevation range creates a long gradient in growing-season temperature. Sites from the western side of Gongga Mountain also tend to be drier than sites at corresponding elevations on the eastern side. By sampling 11 sites over a range of nearly 3000 m in elevation, from both the western and eastern sides, we obtained a data set of hydraulic and photosynthetic traits (see Methods) on woody plants encompassing a wide range of climates.

**Fig. 1.**
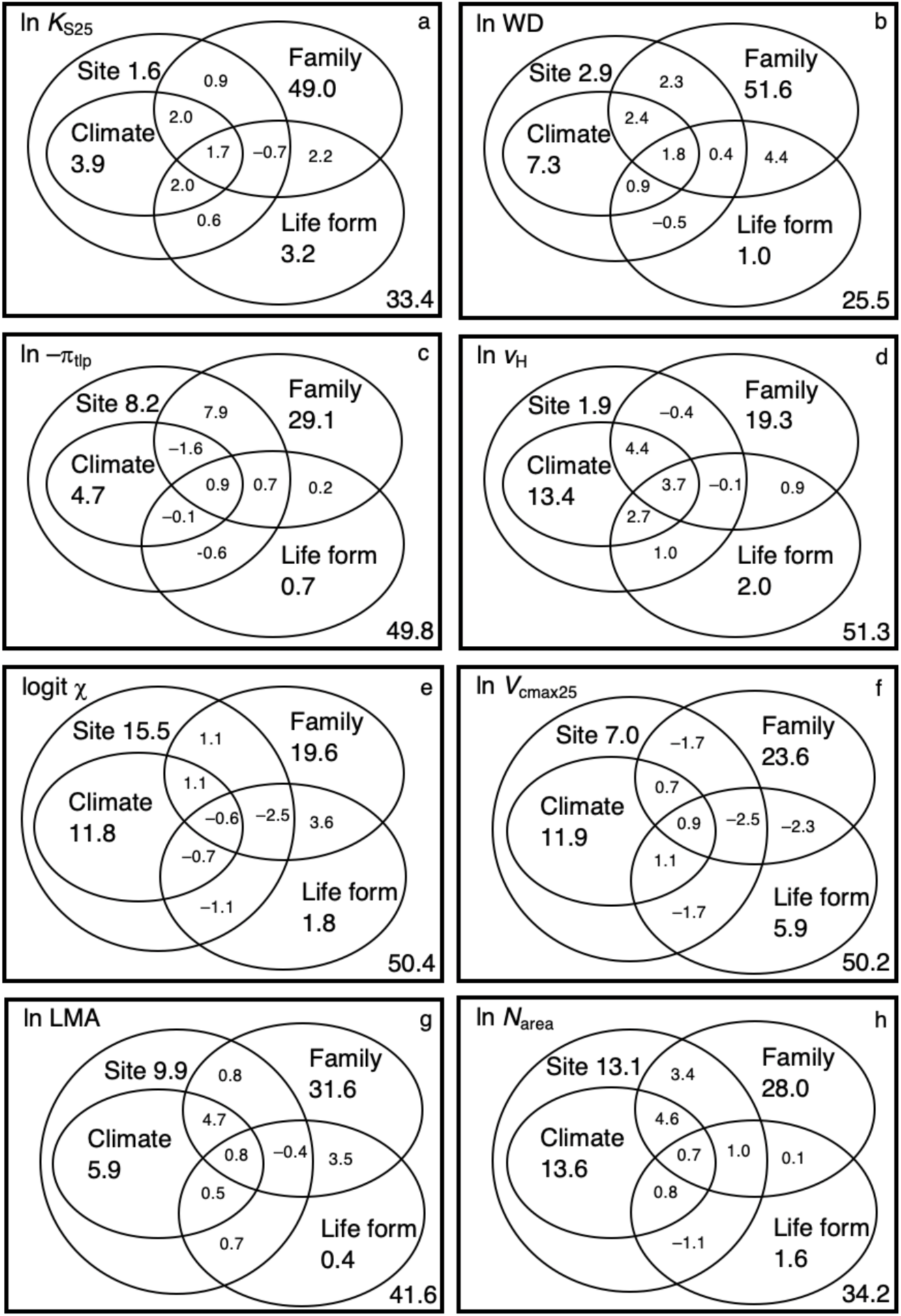
Variance partitioning (%) for each trait. WD is wood density, *K*_S25_ is sapwood-specific hydraulic conductance at 25 °C, π_tlp_ is leaf water potential at turgor loss point, *v*_H_ is the ratio of sapwood to leaf area, LMA is leaf mass per area, χ is the ratio of leaf-internal to ambient CO_2_ partial pressure, *N*_area_ is leaf nitrogen content per area, *V*_cmax25_ is the maximum capacity of carboxylation at 25 °C.

The measured traits can be ranked by phylogenetic influence, according to the fraction of variation explained by family alone in a variation partitioning analysis (Fig. 1). The hydraulic traits wood density (WD) and sapwood-specific hydraulic conductance at 25 °C (*K*_S25_) were most influenced by phylogeny (49-52%); LMA, leaf nitrogen per unit area (*N*_area_) and π_tlp_ were intermediate (28-31%); photosynthetic traits (χ and *V*_cmax_ at 25°C, *V*_cmax25_) and *v*_H_ were least influenced by phylogeny (19-24%). These rankings are approximately mirrored by the percentages of variation explained by site factors and climate (Fig. 1).

Path analysis (Fig. 2) was used to test a framework for trait coordination, based on the hypothesis that the traits that are structurally dependent and more phylogenetically influenced, impose a constraint on more plastic traits, with *v*_H_ as the key trait linking the two sets of traits. Analyses conducted separately on evergreen and deciduous woody plants revealed several general patterns. First, *v*_H_ decreased with *K*_S25_, but increased with *V*_cmax25_ (especially in evergreen plants) (Fig. 2). *K*_S25_ was also lower, and *v*_H_ higher, in plants with high LMA. The leaf economics spectrum (from low to high LMA: Wright, et al. ^20^ thus also influenced *v*_H_, both directly, and indirectly through *K*_S25_. Second, WD was negatively related to *K*_S25_ (especially in deciduous plants), and positively related to –π_tlp_. Third, both LMA and –π_tlp_ negatively influenced χ. In other words, plants with low (more negative) turgor loss point and/or high LMA tend to operate with low χ, with low χ in turn being linked to higher *V*_cmax_ and therefore higher *v*_H_. Fourth, *N*_area_ was found to depend jointly on LMA and *V*_cmax25_, consistent with accumulating evidence – e.g. Dong, et al. ^25^, Xu, et al. ^26^ – for the dependence of *N*_area_ on leaf structure (with LMA as the dominant control, here as in other analyses), and a weaker relationship to *V*_cmax25_. Together, through direct and indirect effects, these hypothesized causal pathways account for all of the significant bivariate relationship among traits (Fig. S2; Table S1).

**Fig. 2.**
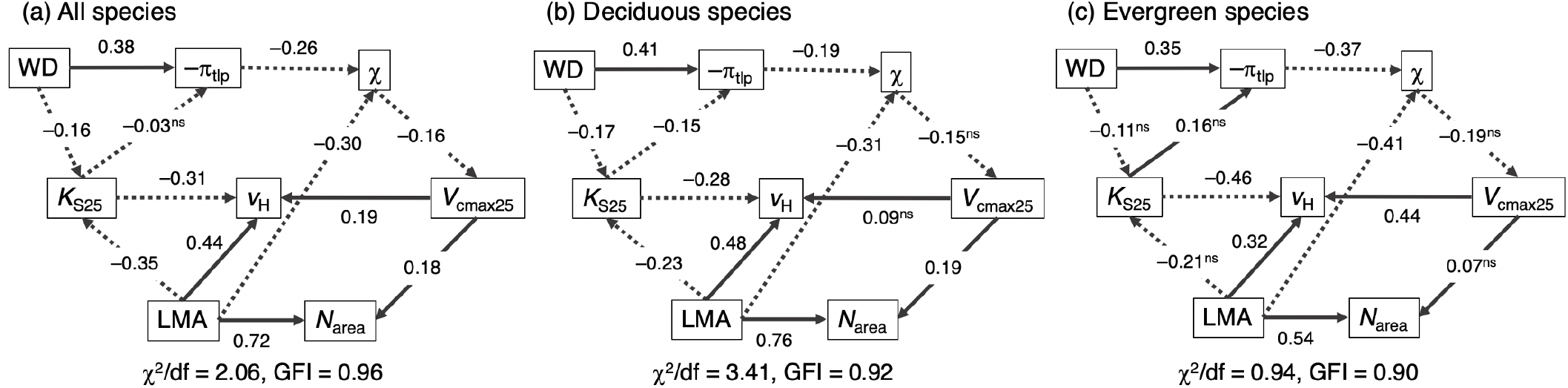
Path analysis of hydraulic and photosynthetic traits for all species (a), separately deciduous (b) and evergreen species (c). WD is wood density, *K*_S25_ is sapwood-specific hydraulic conductance at 25 °C, π_tlp_ is leaf water potential at turgor loss point, *v*_H_ is the ratio of sapwood to leaf area, LMA is leaf mass per area, χ is the ratio of leaf-internal to ambient CO_2_ partial pressure, *N*_area_ is leaf nitrogen content per area, *V*_cmax25_ is the maximum capacity of carboxylation at 25 °C. The arrows indicate the proposed links between traits. Solid lines indicate positive relationships, dotted lines negative relationships. Standard path coefficients are shown near the line (not significant: ns). The trait coordination structure was evaluated using the ratio of χ^2^ and degree of freedom (χ^2^/df) and goodness-of-fit index (GFI).

Our analyses (Figs 2, S2) indicated only a weak trade-off between leaf drought tolerance and xylem hydraulic efficiency. *K*_S25_ and –π_tlp_ were negatively related for deciduous species, but this relationship was not significant for evergreen species, or across all species considered together. The coordination of xylem water transport and stomatal water loss implies that plants should optimally allocate resources so that maximum water transport matches maximum photosynthesis (see Methods for derivations):

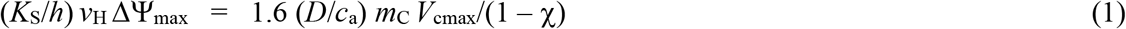

where *h* is the length of the water transport pathway (roughly equivalent to plant height), ΔΨ_max_ is the maximum decrease in water potential from soil to leaves, *D* is the vapour pressure deficit, *c*_a_ is the ambient partial pressure of CO_2_ (dependent on atmospheric pressure), and *m*_C_ = (χ*c*_a_ – Γ^*^)/(χ*c*_a_ + *K*), where Γ^*^ is the photorespiratory compensation point and *K* is the effective Michaelis constant of Rubisco (both dependent on temperature and atmospheric pressure). The factor *m*_C_ reduces photosynthesis under natural conditions relative to *V*_cmax_. In practice, however, *K*_S_ and *h* are not separable because the tapering of xylem elements implies a positive correlation between them that strongly mitigates the effect of path length on whole-stem conductance ^8,9,27^. To test equation (1) we took –π_tlp_ as a surrogate for ΔΨ_max_ ^28^ and subsumed the effect of height in a composite constant, leading to the following relationship:

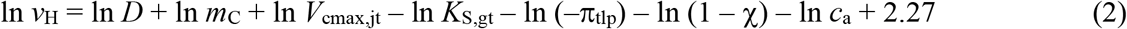

where *V*_cmax,jt_ refers to *V*_cmax_ adjusted to mean July maximum temperature and *K*_S,gt_ refers to *K*_S_ adjusted to the growing-season temperature (see Methods). This theoretical model, including just a single fitted parameter across all species (the intercept, reflecting the effect of height), captures the essential trade-off between *v*_H_ and *K*_S_. Both quantities *v*_H_ and *K*_S_ vary greatly among species (variance of log-transformed variables = 0.4 and 0.73, respectively, averaged across the deciduous and evergreen species-sets; Table S1), allowing a wide variety of hydraulic strategies to coexist within communities. *V*_cmax25_ also varies widely among species (0.69), and more so than either χ (0.28) or π_tlp_ (0.03) (Table S1). The model also predicts a tendency for plants with high *V*_cmax_ to have large *v*_H_, and/or *K*_S_, to allow a correspondingly high rate of water loss. Relationships between *A*_sat_ and plant hydraulic traits found in many studies ^6,29^ are consistent with this prediction. Moreover, equation (2) predicts environmental modulation of the relationship between *v*_H_ and other traits. Specifically, it predicts a positive impact of vapour pressure deficit (*D*) on *v*_H_. As *D* increases, plants are thus expected to allocate relatively less carbon to leaves, and more to stems and roots, resulting in increasing *v*_H_.

Predicted *v*_H_ captured 90% of the observed variation in *v*_H_ across sites (Fig. 3). These predictions (see equation 2), were based on observed hydraulic traits and *c*_a_, and on predicted optimal values of *V*_cmax_, *m*_C_ and χ. Analysis of the modelled contribution of individual factors showed that *K*_S_ was the most important predictor of the variation in site-mean *v*_H_ along the elevation gradient (Fig. 4). The improvement in *R*^2^ contributions for the relationships of predicted ln(*v*_H_) to contributions due to different predictors was 0.42 for *K*_S_, 0.17 for (1 – χ), 0.12 for *c*_a_, 0.11 for *m*_C_, 0.09 for *D*, and 0.05 for *V*_cmax_ and 0.03 for π_tlp_ (Fig. 4a); or in an alternative breakdown of controls, 0.42 for *K*_S_, 0.21 for temperature, 0.17 for radiation, 0.10 for *D*, 0.06 for elevation and 0.03 for π_tlp_ (Fig. 4b).

**Fig.3.**
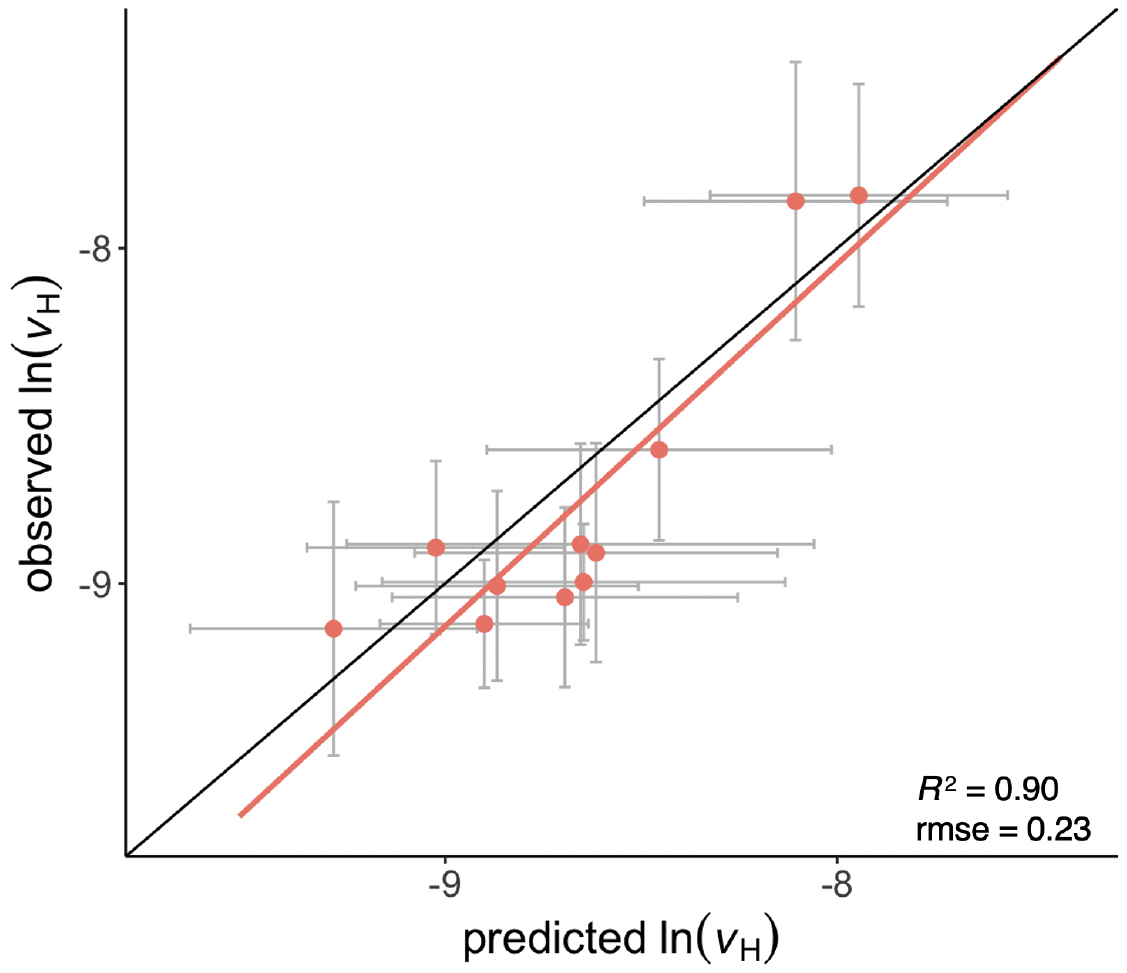
Comparison between site-mean observed and predicted ratios of sapwood to leaf area (v_H_). The error bar on the *y*-axis is the standard deviation of observed ln (*v*_H_) at each site; that on the *x*-axis is the standard deviation of predictions, considering observed variations of sapwood-specific hydraulic conductance (*K*_S_) and leaf water potential at turgor loss point (π_tlp_).

**Fig.4.**
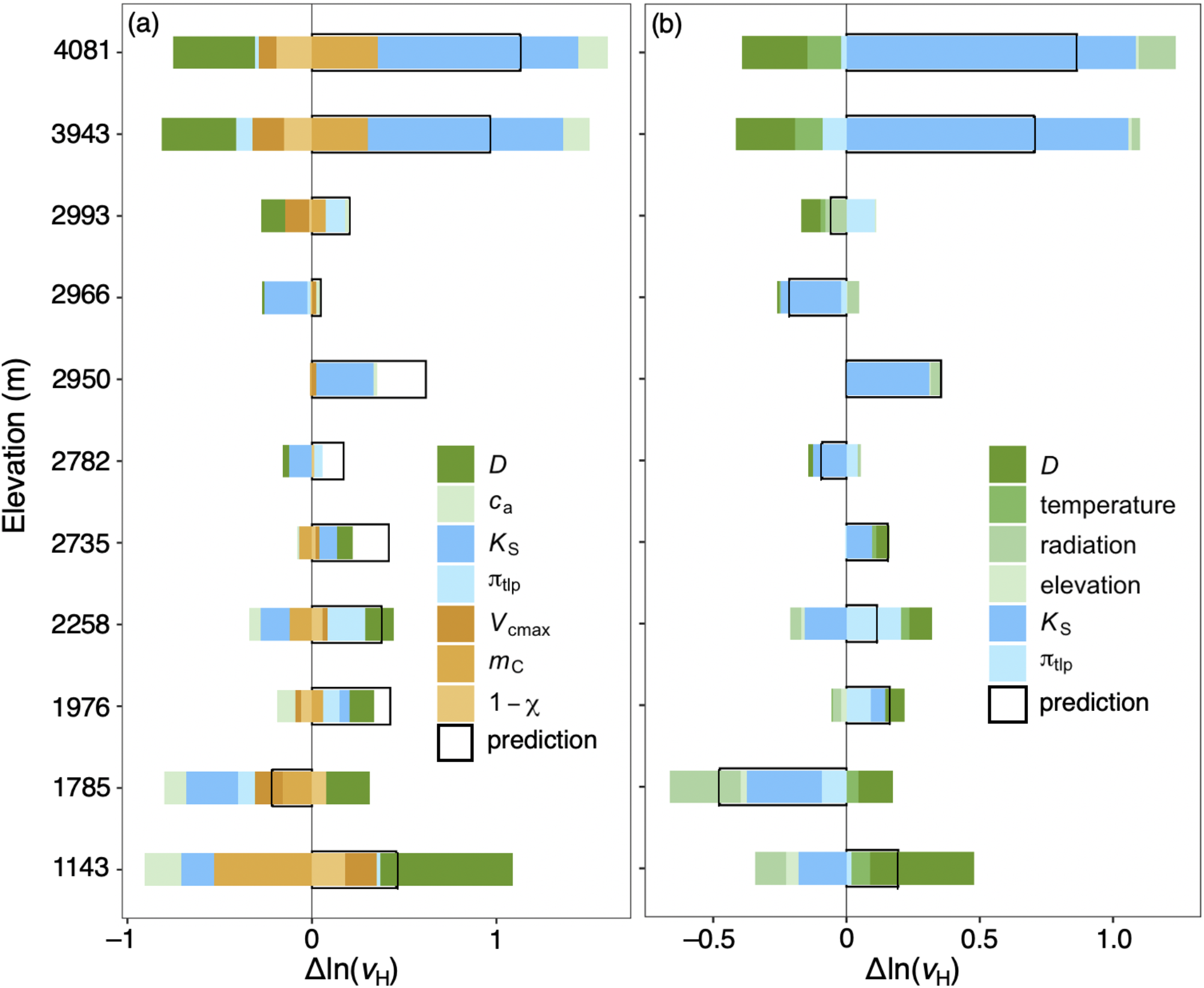
The modelled contribution of different predictors to v_H_ variation at 11 sites sampled along an elevational gradient in the Gongga Mountains, China. (a) The contribution of direct predictors from equation (2). (b) The total contribution of environmental predictors through maximum capacity of carboxylation (*V*_cmax_), the ratio of leaf-internal to ambient CO_2_ partial pressure (χ), *m*_C_ and *c*_a_, along with hydraulic traits. In each panel, the vertical black line represents the baseline ln(*v*_H_) across sites (on which the data were centred, such that the *x*-axis represents the contribution of predictors: Δln(*v*_H_)). Transparent bars with black borders show the changes in predicted values compared to the baseline ln(*v*_H_). Environmental effects are shown in green; photosynthesis-related effects in orange; hydraulic trait effects – sapwood-specific hydraulic conductance (*K*_S_) and leaf water potential at turgor loss point (π_tlp_) – are shown in blue bars.

## Discussion

The results of path analysis (Fig. 2) and the success of the optimality model (Fig. 3) are consistent with the proposed central role of *v*_H_ in coordinating hydraulic and photosynthetic traits ^12^. The *v*_H_ variation mainly results from that in *K*_S_ and χ or temperature. Species deploying a larger total leaf area at a given sapwood area (lower *v*_H_) tend to have higher *K*_S_ ^15^. However, the weak trade-off between π_tlp_ and *K*_S_ implies that low hydraulic safety does not always accompany high *K*_S_. Although the xylem tension at which 50% of the maximum conductivity is lost (*P*_50_) is the most commonly used index of hydraulic safety, we used π_tlp_ for this purpose, noting that the two measures are strongly correlated ^6,30^. Globally, a weak trade-off between hydraulic safety and efficiency has been reported ^31^, although new work suggests a tight trade-off between efficiency and safety may be a feature of climates with highly seasonal precipitation ^32^. That is, plants in environments with less seasonal precipitation need not have high hydraulic efficiency, which may be accompanied by unknown costs or risks.

The relatively rapid acclimation of photosynthetic traits to the local environment ^33^ that leads to the indirect relationship between *v*_H_ and χ (Fig. S2h) has been noted before ^13^. The key role of *v*_H_ in mediating leaf physiology and hydraulics arises because of its plasticity. Variance partitioning (Fig. 1) showed that WD and *K*_S_ are far more strongly linked to phylogeny than other traits. This is presumably because both are related to wood anatomy. *K*_S_ is proportional to the fourth power of mean xylem conduit diameter (according to the Hagen–Poiseuille equation: Tyree and Ewers ^34^), while WD is largely dependent on fibre wall and lumen fractions ^35^. Thus, it might be expected that these traits would show a strong evolutionary convergence within lineages. By contrast, π_tlp_ is known to change after drought through osmotic adjustment ^36^, implying a higher degree of plasticity consistent with the lower influence of family, and the higher influence of environmental factors, on this trait compared to other hydraulic traits (Fig. 1). Plants can adjust *v*_H_ more quickly than other traits related to plant hydraulics – by shedding leaves to balance water supply and demand ^2^. Since photosynthetic traits, particularly χ and *V*_cmax_, respond to environmental conditions on timescales of weeks to months by regulating intrinsic biochemical characteristics ^33,37^, the plasticity of *v*_H_ is required to match photosynthetic acclimation, avoid unnecessary carbon costs, and help to ensure survival under unfavourable conditions.

Wood density has been considered as a crucial trait in a “wood economics spectrum” linking water transport, mechanical support and tree mortality ^38^. Dense wood, found in many species from arid habitats, is generally associated with narrow conduits ^39^ that restrict hydraulic conductance ^40^ but also confers resistance to embolism ^41^, possibly owing to thicker conduit walls and smaller pores in the pit membranes ^42,43^. Wood xylem is the foundation for water transportation, but leaves are often a major bottleneck for water flow, contributing 30% of whole-plant hydraulic resistance on average ^44^. Leaves with lower π_tlp_ can keep their stomata open and continue photosynthesizing at more negative water potentials; on the other hand, this strategy may incur a greater carbon cost to maintain leaf turgor^45^.

The tight relationships among LMA, *v*_H_ and *K*_S_ indicate biologically-important interactions between carbon investment strategy and hydraulics. Leaves with low LMA tend to display a larger leaf area to fix carbon within a relatively short leaf life span. Meanwhile, high hydraulic conductance at both leaf and stem levels ensures that large amount of water can be transported to leaves for transpiration, in order to maintain open stomata and a high rate of CO_2_ uptake ^8,30^. This relationship between hydraulics and LMA may also be associated with physiological characteristics. Thicker leaves (high LMA) tend to have a longer diffusional pathways in the mesophyll, which increases water movement resistance outside leaf xylem and decreases hydraulic conductance ^46^.

The prediction of site-mean *v*_H_ using optimality theory offers a promising approach to implement hydraulics into vegetation and land-surface models. If the focus is on “typical” vegetation in a given climate, the relationship we have predicted (and demonstrated) that applies to site-mean *v*_H_ (equation (2)), with photosynthetic traits predicted by optimality theory, could provide a straightforward way to couple photosynthetic and hydraulic traits in models. However, there is considerable diversity in hydraulic traits (notably *K*_S_) that is linked to LMA, which raises two practical issues: first, how to predict environmental influences on the leaf economics spectrum; second, how to deal with the large within-community variation in both LMA and *K*_S_. Xu, et al. ^26^ have demonstrated a method to predict optimal LMA for deciduous plants, but a different approach is required for evergreen plants ^47^. A way needs to be found to simultaneously estimate the *distribution* of values for highly variable, non-plastic traits. A solution to this problem would be a major step forward for modelling the terrestrial carbon, water and nitrogen cycles.

## Methods

### Model derivation

The theory of *v*_H_ variation is based on Prentice, et al. ^17^. According to Fick’s law and Darcy’s law respectively ^48,49^, transpiration can be calculated from either water demand or supply:

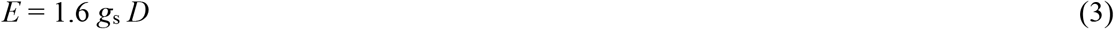

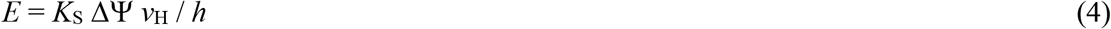

where *E* is the transpiration rate (mol m^−2^ s^−1^), *g*_s_ is stomatal conductance (mol m^−2^ s^−1^), and *D* is the vapour pressure deficit (Pa). Here *h* is the path length (m), roughly equal to plant height; *K*_S_ is the sapwood-specific hydraulic conductance (kg m^−2^ s^−1^ Pa^−1^); *v*_H_ is the ratio of sapwood to leaf area (m^3^ m^−3^); and ΔΨ is the difference between leaf and soil water potential (Ψ_min_ and Ψ_soil_, Pa). We assume ΔΨ has a maximum value equal to –π_tlp_ (Ψ_soil_ ≈ 0 under well-watered conditions) since π_tlp_ is an appropriate proxy for Ψ_min 28_. The uncertainty of the π_tlp_ proxy has little impact on our results as it is not a principal predictor in our model.

From the diffusion equation and the photosynthesis model of Farquhar, von Caemmerer and Berry ^50^, we can calculate *g*_s_ from *V*_cmax_, χ and *m*_C_:

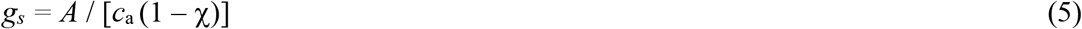

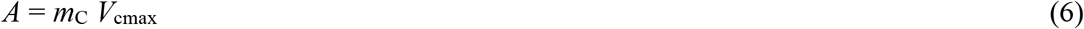

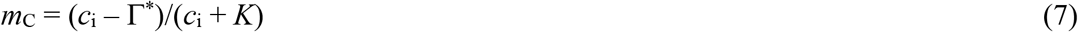

where *A* is the assimilation (photosynthesis) rate (mol m^−2^ s^−1^), *c*_a_ is the ambient partial pressure of CO_2_ (Pa), χ is the ratio of leaf-internal to ambient CO_2_ partial pressure (Pa Pa^−1^), *V*_cmax_ is the maximum capacity of carboxylation (mol m^−2^ s^−1^), Γ_*_ is the photorespiratory compensation point (Pa), and *K* is the effective Michaelis-Menten coefficient of Rubisco (Pa). Substituting *g*_s_ from equations (5) to (7) into equation (3) yields equation (1).

Since *V*_cmax_ acclimates to the environment on a weekly to monthly timescale while *K*_S_ is determined by xylem structure and therefore less able to vary seasonally, we work with *V*_cmax_ at the mean daily maximum temperature in July (*V*_cmax,jt_) and *K*_S_ at the mean daily maximum temperature during the growing season (defined as the period with daytime temperatures > 0 °C) (*K*_S,gt_).

χ in equation (5) can be estimated as follows, based on the least-cost hypothesis ^18^:

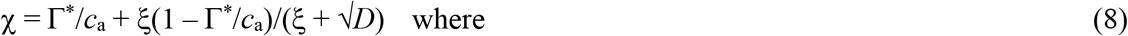

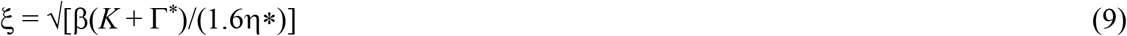

where β is the ratio at 25 °C of the unit costs of maintaining carboxylation and transpiration capacities (146, based on a global compilation of leaf δ^13^C measurements), and η* is the viscosity of water relative to its value at 25 °C.

*V*_cmax,jt_ can also be predicted from climate ^19^:

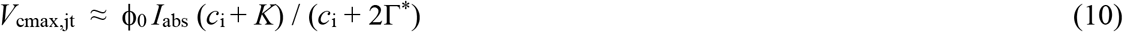

where ϕ_0_ is the intrinsic quantum efficiency of photosynthesis (which we assign the value 0.085 μmol C μmol^−1^ photon), *I*_abs_ is the photosynthetic photon flux density (PPFD) absorbed by leaves (mol m^−2^ s^−1^), and *c*_i_ is the leaf-internal CO_2_ partial pressure (*c*_i_ = χ *c*_a_) (Pa).

### Trait data

Trait data were measured at 11 sites in late July 2018 and August 2019, during the active growing season, in the Gongga Mountain region (29° 34’ 16” – 29° 54’ 52” N and 101° 59’ 08” – 102° 9’ 42” E, Fig. S1). We collected the data needed to allow the calculation of four leaf traits: leaf mass per area (LMA), leaf nitrogen per unit area (*N*_area_), the maximum capacity of carboxylation (*V*_cmax_), and the ratio of leaf-internal to ambient CO_2_ partial pressure (χ). Hydraulic traits, specifically the ratio of sapwood to leaf area (Huber value, *v*_H_), sapwood-specific hydraulic conductance (*K*_S_), wood density (WD) and leaf potential at turgor loss point (π_tlp_), were measured on all the woody broad-leaved species. We sampled all the tree species and at least five shrub species at each site. All samples were taken from the outer and upper canopy receiving direct sunshine.

LMA was calculated from the measurements of leaf area and dry weight following standard protocols ^51^. Multiple leaves, or leaflets for compound leaves, were randomly selected and scanned using a Canon LiDE 220 Scanner. The area was estimated in Matlab. The dry weight of these leaves was measured after oven-drying at 75 °C for 48 h to constant weight. We calculated LMA as the ratio of dry mass to leaf area. Leaf nitrogen content was measured using an Isotope Ratio Mass Spectrometer (Thermo Fisher Scientific Inc., USA). *N*_area_ was calculated from LMA and leaf nitrogen content. The LMA value for a species at a given site was the average of three separate measurements made on leaves from multiple individuals, while *N*_area_ measurements were made on pooled samples of leaves from multiple individuals.

Carbon isotopic values (δ^13^C) were measured using an Isotope Ratio Mass Spectrometer (Thermo Fisher Scientific Inc., USA). Values were measured on pooled samples of leaves from multiple individuals. Estimates of χ were made using the method of Cornwell, et al. ^52^ to calculate isotopic discrimination (Δ) from δ^13^C with a standard formula using the recommended values of *a’* and *b’* of 4.4 ‰ and 27 ‰, respectively ^53,54^:

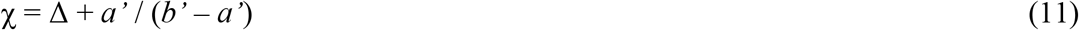

Leaf gas-exchange measurements were made in the field using a portable infrared gas analyser (IRGA) system (LI-6400; Li-Cor Inc., Lincoln, NB, USA). Sunlit branches from the outer canopy were collected and re-cut under water immediately prior to measurement. In-situ measurements were taken with relative humidity and chamber block temperature similar to the ambient conditions, and a constant airflow rate (500 μmol s^−1^). *V*_cmax_ at leaf temperature (*V*_cmax,lt_) was calculated from the light-saturated rate of net CO_2_ fixation at ambient CO_2_, measured on one individual of each species, using the one-point method ^55^ and adjusted to a standard temperature of 25 °C (*V*_cmax25_) and maximum temperature in July (*V*_cmax,jt_) using the method of Bernacchi, et al. ^56^.

Branches with a diameter wider than 7 mm were sampled for hydraulic traits. We measured the cross-sectional area of the xylem at both ends of each cut branch using digital calipers. Sapwood area was calculated as the average of these two measurements. All leaves attached to the branch were removed and dried in an oven at 70 °C for 72 hours before weighing. The total leaf area was obtained from dry mass and LMA. The ratio of sapwood area and leaf area was calculated as *v*_H_. The *v*_H_ value of one species at each site was the average of three measurements made on branches from different individuals.

Five branches from at least three mature individuals of the same species at each site were collected, wrapped in moist towels and sealed in black plastic bags, and then immediately transported to the laboratory. All the samples were re-cut under water, put into water and sealed in black plastic bags to rehydrate overnight. *K*_S_ was measured using the method described in Sperry, et al. ^57^. Segments (10 - 15 cm length) were cut from the rehydrated branches and flushed using 20 mmol L^−1^ KCl solution for at least 30 minutes (to remove air from the vessels) until constant fluid dripped from the segment section. The segments were then placed under 0.005 MPa pressure to record the time (*t*) they took to transport a known water volume (*W*, m^3^). Length (*L*, m), sapwood areas of both ends (*S*_1_ and *S*_2_, m^2^) and temperature (*T*_m_, °C) were recorded. Sapwood-specific hydraulic conductance at measurement temperature (*K*_S,m_ kg m^−2^ s^−1^ MPa^−1^) was calculated using equation (12). This was transformed to *K*_S_ at mean maximum temperature during the growing season (*K*_S,gt_) and standard temperature (*K*_S25_) following equations (13) and (14):

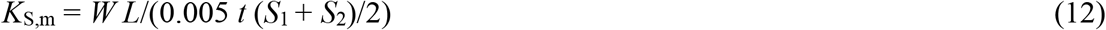

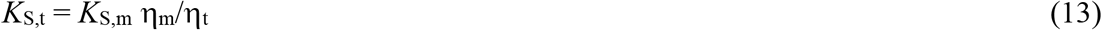

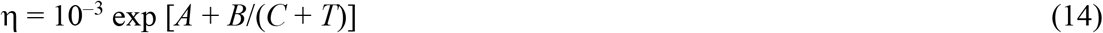

where η_m_ and η_t_ are the water viscosity at measurement temperature and transformed temperature (i.e. mean maximum daytime temperature during the growing season and standard temperature, 25 °C in this study), respectively. The parameter values adopted in (14) were *A* = −3.719, *B* = 580 and *C* = −138^58^.

A small part of each sapwood segment was used to measure wood density, the ratio of dry weight to volume of sapwood. After removal of bark and heartwood, the displacement method was used to measure the volume of sapwood and the dry weight of sapwood was obtained after oven-drying at 70 °C for 72 hours to constant weight. Wood density was calculated as the ratio of the volume and the dry weight of sapwood.

We applied the method described by Bartlett, et al. ^45^ for the rapid determination of π_tlp_. After rehydration overnight, discs were sampled from mature, healthy leaves collected on each branch, avoiding major and minor veins and using a 6-mm-diameter puncher. Leaf discs wrapped in foil were frozen in liquid nitrogen for at least 2 minutes and then punctured 20 times quickly with sharp-tipped tweezers. Five repeat experiments using leaves from multiple individuals were carried out for every species at each site. We measured osmotic potential (π_osm_) with a VAPRO 5600 vapor pressure osmometer (Wescor, Logan, UT, USA) and calculated π_tlp_ (in MPa) as:

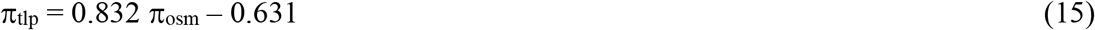

### Climate data

We derived climate variables at each of the 11 sampled sites using meteorological data (monthly maximum and minimum temperature, fraction of sunshine hours and water vapour pressure) from 17 weather stations in the Gongga region: (http://data.cma.cn/data/cdcdetail/dataCode/SURF_CLI_CHN_MUL_MON.html) and the elevationally-sensitive ANUSPLIN interpolation scheme^59^. The meteorological data were available from January 2017 to December 2019. The monthly data were converted to daily values by linear interpolation in order to calculate the bioclimatic variables mean maximum temperature during the growing-season (defined as the period with daytime temperature > 0 °C), growing-season mean PPFD, and vapour pressure deficit under maximum daytime temperature in July, using the Simple Process-Led Algorithms for Simulating Habitats (SPLASH) model ^60^.

### Data analysis

All statistical analyses were carried out in R3.1.3 (R Core Team 2015). To homogenize the variance, traits were natural log-transformed and χ was logit-transformed; for π_tlp_, the absolute value (–π_tlp_) was natural log-transformed. Trait variance partitioning was carried out using the *vegan* package _61_. Path analysis was used to characterize the trait coordination framework built on the idea that plastic traits are influenced by structural traits, using the *lavaan* package ^62^. The model was evaluated using the ratio of χ^2^ and degree of freedom (χ^2^/df) and goodness-of-fit index (GFI). The χ^2^/df of models using all species and only evergreen species were below 3, and the GFI of all three models were larger than 0.9 (Fig. 2). Traits under standard conditions (*V*_cmax25_ and *K*_S25_) were used in variance partitioning, path analysis and bivariate regressions to eliminate the effect of temperature, while trait values under growth conditions were used for theoretical prediction.

To examine the importance of each predictor in equation (2) for the prediction of *v*_H_, we evaluated the contributions of each variable in four steps as follows. First we evaluated the contributions of predictors included in the equation (*D, m*_C_, *V*_cmax_, *K*_S_, π_tlp_, *c*_a_, 1 – χ), and also the total contributions from indirect predictors driving photosynthesis-related traits (*D*, temperature, radiation, elevation) along with the hydraulic predictors. The baseline value of each predictor was the median value of its site-mean values across the 11 sites. These baseline values were used to generate baseline predicted ln(*v*_H_). Second, one predictor after another was changed to actual values at each site, while other predictors remained at their baseline values. We then calculated ln(*v*_H_) using these inputs. Third, the contribution of each predictor at each site was calculated as the difference between values from the second and first step (indexed as Δln(*v*_H_) in Fig. 4). Last, we calculated the improvement in *R*^2^ of the relationships between predicted ln(*v*_H_) and contributions of each predictor across sites. *R*^2^ improvements due to each variable were averaged over orderings among predictors, yielding the relative importance of each variable. This procedure was run using the *relaimpo* package ^63^.

## Data availability

The trait data will be released at time of publication.

## Acknowledgements

We thank Yingying Ji, Meng Li, Xinyu Liu, Giulia Mengoli, Yunke Peng, Shengchao Qiao, Yifan Su, Yuhui Wu, Shuxia Zhu and Wei Zheng for their assistance in collecting trait data in Gongga Mountain. We also thank Zonghan Ma for help with interpolation of the climate data. This work was funded by National Science Foundation China (grant no. 31971495, 32022052, 91837312). I.C.P. and S.P.H are supported by the High-End Foreign Expert program of the China State Administration of Foreign Expert Affairs at Tsinghua University (GDW20191100161) I.C.P. acknowledges support from the European Research Council (787203 REALM) under the European Union’s Horizon 2020 research programme. S.P.H. is supported by the European Research Council (694481 GC2.0) under the same programme. I.J.W. acknowledges support from the Australian Research Council (DP170103410).

## Author Contributions

H.X. carried out the analyses and prepared the manuscript with contributions from all co-authors. H.W., S.P.H. and I.C.P. designed the fieldwork, collected samples and measured plant traits. I.C.P and I.J.W developed and extended the least-cost theory. All authors contributed to the interpretation of the results.

## References

1. Rowland, L. et al. Death from drought in tropical forests is triggered by hydraulics not carbon starvation. Nature 528, 119–122, doi:10.1038/nature15539 (2015).

2. Choat, B. et al. Triggers of tree mortality under drought. Nature 558, 531–539, doi:10.1038/s41586-018-0240-x (2018).

3. Brodribb, T. J. Xylem hydraulic physiology: The functional backbone of terrestrial plant productivity. Plant Science 177, 245–251, doi:10.1016/j.plantsci.2009.06.001 (2009).

4. Scoffoni, C. et al. Hydraulic basis for the evolution of photosynthetic productivity. Nature Plants 2, 16072, doi:10.1038/nplants.2016.72 (2016).

5. Brodribb, T. J., Feild, T. S. & Jordan, G. J. Leaf maximum photosynthetic rate and venation are linked by hydraulics. Plant Physiology 144, 1890–1898, doi:10.1104/pp.107.101352 (2007).

6. Zhu, S. D. et al. Leaf turgor loss point is correlated with drought tolerance and leaf carbon economics traits. Tree Physiology 38, 658–663, doi:10.1093/treephys/tpy013 (2018).

7. Deans, R. M., Brodribb, T. J., Busch, F. A. & Farquhar, G. D. Optimization can provide the fundamental link between leaf photosynthesis, gas exchange and water relations. Nature Plants 6, 1116–1125, doi:10.1038/s41477-020-00760-6 (2020).

8. Mencuccini, M. et al. Leaf economics and plant hydraulics drive leaf : wood area ratios. New Phytologist 224, 1544–1556, doi:10.1111/nph.15998 (2019).

9. Christoffersen, B. O. et al. Linking hydraulic traits to tropical forest function in a size-structured and trait-driven model (TFS v.1-Hydro). Geoscientific Model Development 9, 4227–4255, doi:10.5194/gmd-9-4227-2016 (2016).

10. Mencuccini, M., Manzoni, S. & Christoffersen, B. Modelling water fluxes in plants: from tissues to biosphere. New Phytologist 222, 1207–1222, doi:10.1111/nph.15681 (2019).

11. De Kauwe, M. G. et al. A test of an optimal stomatal conductance scheme within the CABLE land surface model. Geoscientific Model Development 8, 431–452, doi:10.5194/gmd-8-431-2015 (2015).

12. Rosas, T. et al. Adjustments and coordination of hydraulic, leaf and stem traits along a water availability gradient. New Phytologist 223, 632–646, doi:10.1111/nph.15684 (2019).

13. Martinez-Vilalta, J. et al. Hydraulic adjustment of Scots pine across Europe. New Phytologist 184, 353–364, doi:10.1111/j.1469-8137.2009.02954.x (2009).

14. Liu, H. et al. Hydraulic traits are coordinated with maximum plant height at the global scale. Science Advances 5, eaav1332 (2019).

15. Togashi, H. F. et al. Morphological and moisture availability controls of the leaf area-to-sapwood area ratio: analysis of measurements on Australian trees. Ecology and Evolution 5, 1263–1270, doi:10.1002/ece3.1344 (2015).

16. Gleason, S. M., Butler, D. W. & Waryszak, P. Shifts in leaf and stem hydraulic traits across aridity gradients in Eastern Australia. International Journal of Plant Sciences 174, 1292–1301, doi:10.1086/673239 (2013).

17. Prentice, I. C., Dong, N., Gleason, S. M., Maire, V. & Wright, I. J. Balancing the costs of carbon gain and water transport: testing a new theoretical framework for plant functional ecology. Ecology Letters 17, 82–91, doi:10.1111/ele.12211 (2014).

18. Wang, H. et al. Towards a universal model for carbon dioxide uptake by plants. Nature Plants 3, 734–741, doi:10.1038/s41477-017-0006-8 (2017).

19. Smith, N. G. et al. Global photosynthetic capacity is optimized to the environment. Ecology Letters 22, 506–517, doi:10.1111/ele.13210 (2019).

20. Wright, I. J. et al. The worldwide leaf economics spectrum. Nature 428, 821 (2004).

21. Poorter, H., Niinemets, U., Poorter, L., Wright, I. J. & Villar, R. Causes and consequences of variation in leaf mass per area (LMA): a meta-analysis. New Phytologist 182, 565–588, doi:10.1111/j.1469-8137.2009.02830.x (2009).

22. Franklin, O. et al. Organizing principles for vegetation dynamics. Nature Plants 6, 444–453, doi:10.1038/s41477-020-0655-x (2020).

23. Lavergne, A., Sandoval, D., Hare, V. J., Graven, H. & Prentice, I. C. Impacts of soil water stress on the acclimated stomatal limitation of photosynthesis: Insights from stable carbon isotope data. Global Change Biology 26, 7158–7172, doi:10.1111/gcb.15364 (2020).

24. Sperry, J. S. et al. Predicting stomatal responses to the environment from the optimization of photosynthetic gain and hydraulic cost. Plant, Cell & Environment 40, 816–830, doi:10.1111/pce.12852 (2017).

25. Dong, N. et al. Leaf nitrogen from first principles: field evidence for adaptive variation with climate. Biogeosciences 14, 481–495, doi:10.5194/bg-14-481-2017 (2017).

26. Xu, H. et al. Predictability of leaf traits with climate and elevation: a case study in Gongga Mountain, China. Tree Physiol, doi:10.1093/treephys/tpab003 (2021).

27. Olson, M. E., Anfodillo, T., Gleason, S. M. & McCulloh, K. A. Tip-to-base xylem conduit widening as an adaptation: causes, consequences, and empirical priorities. New Phytologist 229, 1877–1893, doi:10.1111/nph.16961 (2021).

28. Hochberg, U., Rockwell, F. E., Holbrook, N. M. & Cochard, H. Iso/anisohydry: A plant-environment interaction rather than a simple hydraulic trait. Trends in Plant Science 23, 112–120, doi:10.1016/j.tplants.2017.11.002 (2018).

29. Santiago, L. S. et al. Leaf photosynthetic traits scale with hydraulic conductivity and wood density in Panamanian forest canopy trees. Oecologia 140, 543–550, doi:10.1007/s00442-004-1624-1 (2004).

30. Joshi, J. et al. Towards a unified theory of plant photosynthesis and hydraulics. bioRxiv, 2020.2012.2017.423132, doi:10.1101/2020.12.17.423132 (2020).

31. Gleason, S. M. et al. Weak tradeoff between xylem safety and xylem-specific hydraulic efficiency across the world’s woody plant species. New Phytologist 209, 123–136, doi:10.1111/nph.13646 (2016).

32. Liu, H., Ye, Q., Gleason, S. M., He, P. & Yin, D. Weak tradeoff between xylem hydraulic efficiency and safety: climatic seasonality matters. New Phytologist, doi:10.1111/nph.16940 (2020).

33. Smith, N. G. & Dukes, J. S. Short-term acclimation to warmer temperatures accelerates leaf carbon exchange processes across plant types. Global Change Biology 13, 4840–4853 (2017).

34. Tyree, M. T. & Ewers, F. W. The hydraulic architecture of trees and other woody plants. New Phytologist 119, 345–360 (1991).

35. Ziemińska, K., Butler, D. W., Gleason, S. M., Wright, I. J. & Westoby, M. Fibre wall and lumen fractions drive wood density variation across 24 Australian angiosperms. AoB PLANTS 5, doi:10.1093/aobpla/plt046 (2013).

36. Bartlett, M. K. et al. Global analysis of plasticity in turgor loss point, a key drought tolerance trait. Ecology Letters 17, 1580–1590, doi:10.1111/ele.12374 (2014).

37. Cavanagh, A. P. & Kubien, D. S. Can phenotypic plasticity in Rubisco performance contribute to photosynthetic acclimation? Photosynthesis Research 119, 203–214 (2014).

38. Chave, J. et al. Towards a worldwide wood economics spectrum. Ecology Letters 12, 351–366, doi:10.1111/j.1461-0248.2009.01285.x (2009).

39. Hacke, U. G. & Sperry, J. S. Functional and ecological xylem anatomy. Perspectives in Plant Ecology, Evolution and Systematics 4, 97–115, doi:10.1078/1433-8319-00017 (2001).

40. Zanne, A. E. et al. Angiosperm wood structure: Global patterns in vessel anatomy and their relation to wood density and potential conductivity. American Journal of Botany 97, 207–215, doi:10.3732/ajb.0900178 (2010).

41. Anderegg, W. R. et al. Meta-analysis reveals that hydraulic traits explain cross-species patterns of drought-induced tree mortality across the globe. Proceedings of the National Academy of Sciences of the United States of America 113, 5024–5029, doi:10.1073/pnas.1525678113 (2016).

42. Hacke, U. G., Sperry, J. S., Pockman, W. T., Davis, S. D. & McCulloh, K. A. Trends in wood density and structure are linked to prevention of xylem implosion by negative pressure. Oecologia 126, 457–461, doi:10.1007/s004420100628 (2001).

43. Pittermann, J. et al. The relationships between xylem safety and hydraulic efficiency in the Cupressaceae: the evolution of pit membrane form and function. Plant Physiology 153, 1919–1931, doi:10.1104/pp.110.158824 (2010).

44. Sack, L. & Holbrook, N. M. Leaf hydraulics. Annual Review of Plant Biology 57, 361–381, doi:10.1146/ (2006).

45. Bartlett, M. K. et al. Rapid determination of comparative drought tolerance traits: using an osmometer to predict turgor loss point. Methods in Ecology and Evolution 3, 880–888, doi:10.1111/j.2041-210X.2012.00230.x (2012).

46. Flexas, J., Scoffoni, C., Gago, J. & Sack, L. Leaf mesophyll conductance and leaf hydraulic conductance: an introduction to their measurement and coordination. Journal of Experimental Botany 64, 3965–3981, doi:10.1093/jxb/ert319 (2013).

47. Wang, H. et al. Leaf economics explained by optimality principles. bioRxiv, 2021.2002.2007.430028, doi:10.1101/2021.02.07.430028 (2021).

48. Fick, A. Ueber Diffusion. Annalen der Physik 170, 59–86, doi:https://doi.org/10.1002/andp.18551700105 (1855).

49. Whitehead, D. Regulation of stomatal conductance and transpiration in forest canopies. Tree Physiology 18, 633–644, doi:10.1093/treephys/18.8-9.633 (1998).

50. Farquhar, G. D., von Caemmerer, S. & Berry, J. A. A biochemical model of photosynthetic CO2 assimilation in leaves of C3 species. Planta 149, 78–90, doi:10.1007/BF00386231 (1980).

51. Cornelissen, J. H. C. et al. A handbook of protocols for standardised and easy measurement of plant functional traits worldwide. Australian Journal of Botany 51, 335–380, doi:10.1071/BT02124 (2003).

52. Cornwell, W. K. et al. Climate and soils together regulate photosynthetic carbon isotope discrimination within C3 plants worldwide. Glob Ecol Biogeogr 27, 1056–1067, doi:10.1111/geb.12764 (2018).

53. Farquhar, G. D., Ehleringer, J. R. & Hubick, K. T. Carbon isotope discrimination and photosynthesis. Annual Review of Plant Biology 40, 503–537 (1989).

54. Cernusak, L. A. et al. Environmental and physiological determinants of carbon isotope discrimination in terrestrial plants. New Phytologist 200, 950–965, doi:10.1111/nph.12423 (2013).

55. De Kauwe, M. G. et al. A test of the ‘one-point method’ for estimating maximum carboxylation capacity from field-measured, light-saturated photosynthesis. New Phytologist 210, 1130–1144, doi:10.1111/nph.13815 (2016).

56. Bernacchi, C. J., Singsaas, E. L., Pimentel, C., Portis Jr, A. R. & Long, S. P. Improved temperature response functions for models of Rubisco-limited photosynthesis. Plant, Cell & Environment 24, 253–259 (2001).

57. Sperry, J. S., Donnelly, J. R. & Tyree, M. T. A method for measuring hydraulic conductivity and embolism in xylem. Plant, Cell & Environment 11, 35–40 (1988).

58. Vogel, H. Temperaturabhängigkeitsgesetz der Viskosität von Flüssigkeiten. Physik Z 22, 645–646 (1921).

59. Hutchinson, M. F. & Xu, T. Anusplin version 4.2 user guide. Centre for Resource and Environmental Studies, The Australian National University, Canberra 54 (2004).

60. Davis, T. W. et al. Simple process-led algorithms for simulating habitats (SPLASH v.1.0): robust indices of radiation, evapotranspiration and plant-available moisture. Geoscientific Model Development 10, 689–708, doi:10.5194/gmd-10-689-2017 (2017).

61. Oksanen, J. et al. vegan: Community Ecology Package. R package version 2.4-4. 2, 1–295 (2017).

62. Rosseel, Y. Lavaan: An R package for structural equation modeling and more. Version 0.5– 12 (BETA). Journal of statistical software 48, 1–36 (2012).

63. Groemping, U. Relative Importance for Linear Regression in R: The Package relaimpo. Journal of Statistical Software 17, 925–933 (2006).

