## Supplemental Tables and Figures for "Plant hydraulics coordinated with photosynthetic traits and climate"

**Table S1 Variance-covariance matrices of traits for deciduous and evergreen species.** The values were calculated across all species. All traits were log_e_-transformed except χ which was logit transformed. Trait abbreviations as follows: wood density (WD), sapwood-specific hydraulic conductance at 25 ˚C (*K*_S25_), leaf water potential at turgor loss point (π_tlp_), the ratio of sapwood to leaf area (*v*_H_), leaf mass per area (LMA), the ratio of leaf-internal to ambient CO_2_ partial pressure (χ), leaf nitrogen content per area (*N*_area_) and maximum capacity of carboxylation at 25 ˚C (*V*_cmax25_). *P* values are indicated “***” (< 0.001), “**” (< 0.01) and “*” (< 0.05).

| Deciduous | *v*_H_ | *K*_S25_ | –π_tlp_ | WD | LMA | *N*_area_ | *V*_cmax25_ | logit χ |
| --- | --- | --- | --- | --- | --- | --- | --- | --- |
| *v*_H_ | 0.36 |  |  |  |  |  |  |  |
| *K*_S25_ | –0.19*** | 0.70 |  |  |  |  |  |  |
| –π_tlp_ | 0.02* | –0.03** | 0.03 |  |  |  |  |  |
| WD | 0.03** |  | 0.02*** | 0.05 |  |  |  |  |
| LMA | 0.11*** | –0.06** | 0.02*** |  | 0.12 |  |  |  |
| *N*_area_ | 0.11*** |  | 0.02** |  | 0.11*** | 0.18 |  |  |
| *V*_cmax25_ |  |  |  |  |  | 0.09*** | 0.58 |  |
| logit χ |  |  | –0.02*** |  | –0.06*** | –0.08*** |  | 0.26 |

| Evergreen | *v*_H_ | *K*_S25_ | –π_tlp_ | WD | LMA | *N*_area_ | *V*_cmax25_ | logit χ |
| --- | --- | --- | --- | --- | --- | --- | --- | --- |
| *v*_H_ | 0.43 |  |  |  |  |  |  |  |
| *K*_S25_ | –0.33** | 0.76 |  |  |  |  |  |  |
| –π_tlp_ |  |  | 0.03 |  |  |  |  |  |
| WD |  |  |  | 0.05 |  |  |  |  |
| LMA | 0.14** |  |  |  | 0.13 |  |  |  |
| *N*_area_ | 0.14** |  |  |  | 0.07** | 0.12 |  |  |
| *V*_cmax25_ | 0.36** |  |  |  |  |  | 0.79 |  |
| logit χ |  |  |  |  | –0.07* | –0.09* |  | 0.30 |

**Fig. S1** **Locations of sampling sites and weather stations.** The green dots are the sampling sites and red triangles are the weather stations. The latitude and longitude for the outer plot are in black, inset in navy blue. The background colour represents the elevation gradient.

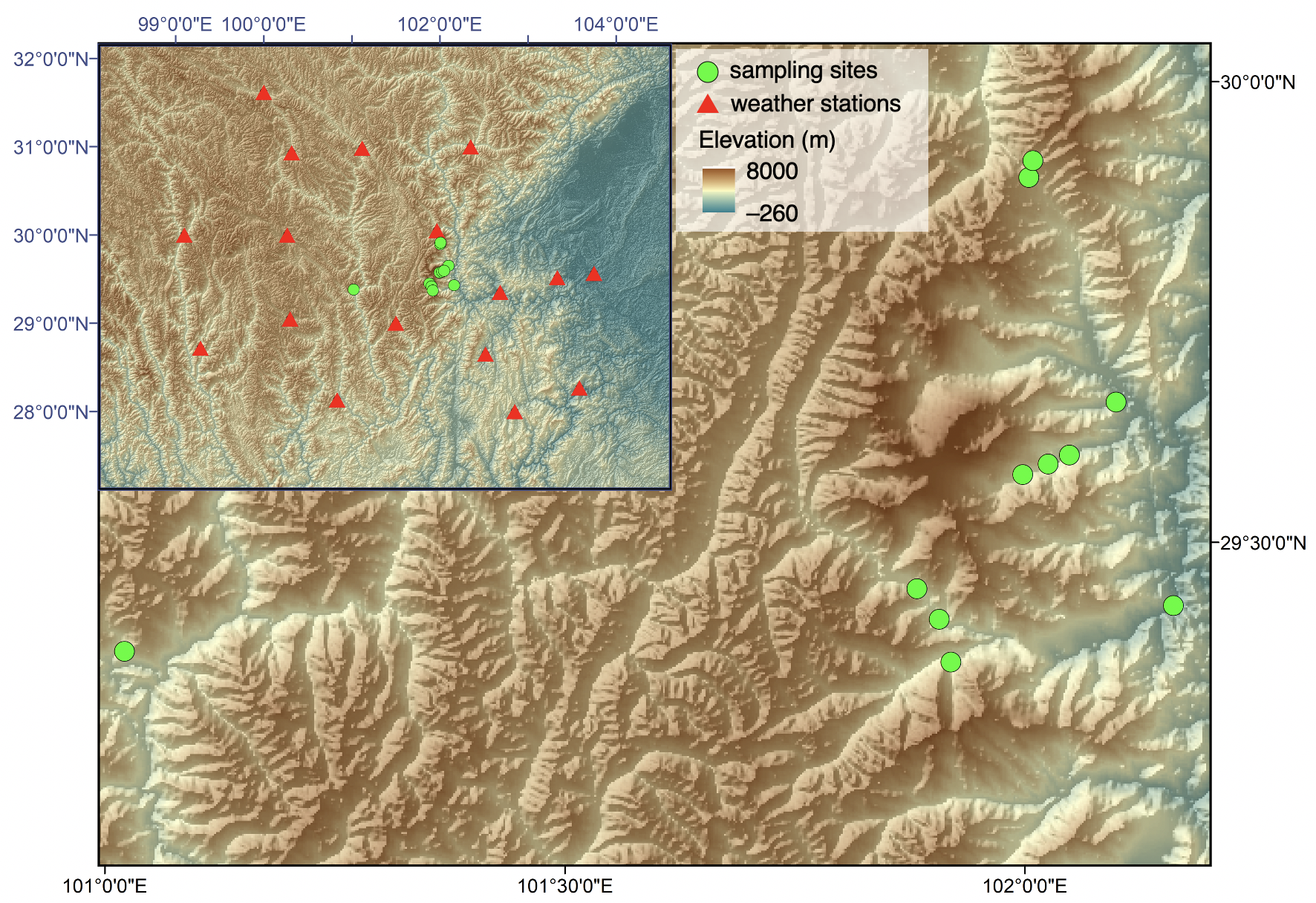

**Fig. S2** **Bivariate relationships between traits.** Deciduous and evergreen species are in blue and red circles, respectively. Solid lines indicate significant relationships (*p* < 0.05), dashed lines nonsignificant relationships. The traits are wood density (WD), sapwood-specific hydraulic conductance at 25 ˚C (*K*_S25_), leaf water potential at turgor loss point (π_tlp_), the ratio of sapwood to leaf area (*v*_H_), leaf mass per area (LMA), the ratio of leaf-internal to ambient CO_2_ partial pressure (χ), leaf nitrogen content per area (*N*_area_) and maximum capacity of carboxylation at 25 ˚C (*V*_cmax25_).

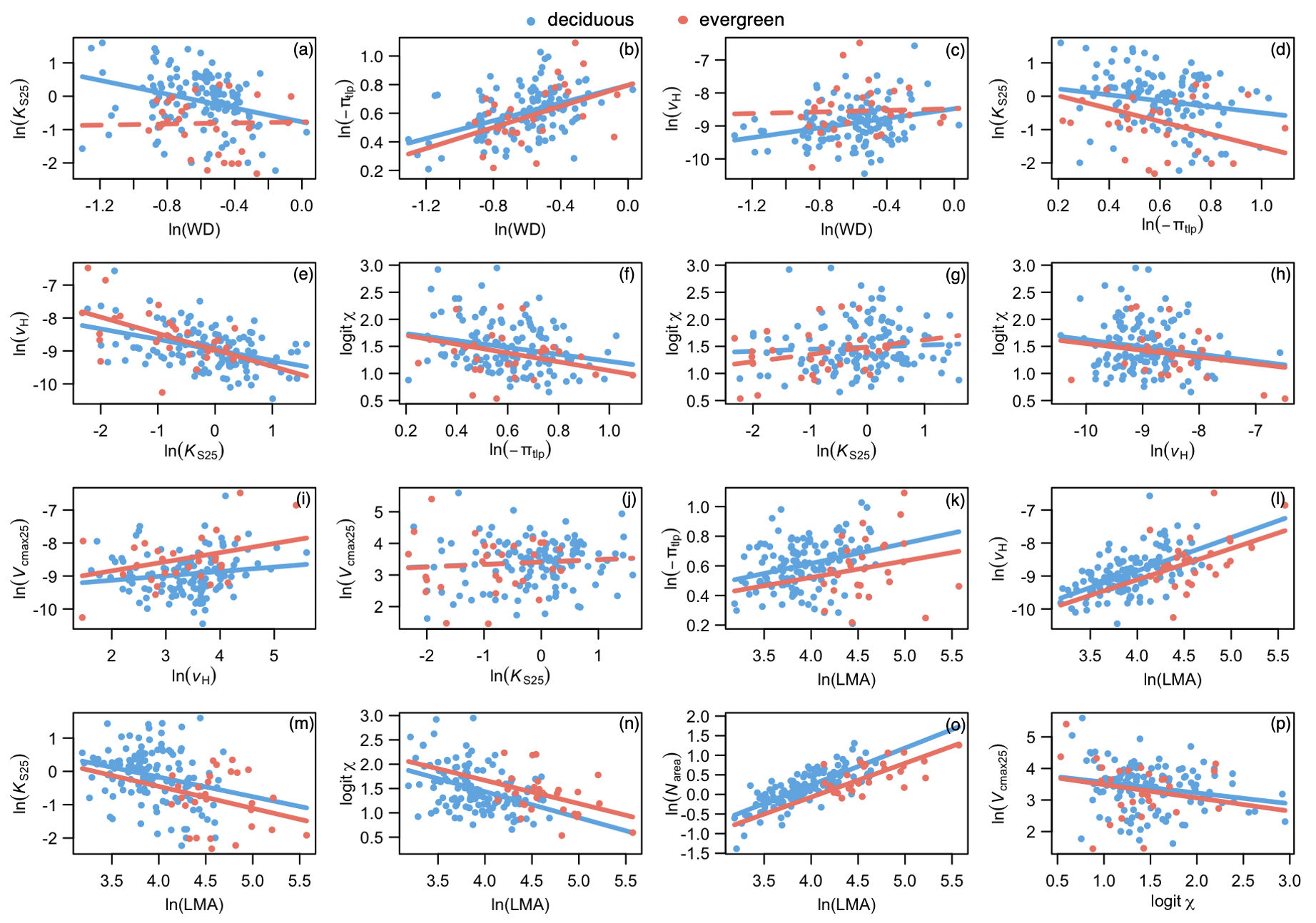
